# Estimand-aware and donor-aware triangulation of predefined gene-set signals across human tendon transcriptomic datasets

**DOI:** 10.64898/2026.09.11.750964

**Authors:** Yushuo Liu, Zitao Wang, Qiuyuan Peng, Yang Li, Bingao Chen

**Affiliations:** School of Physical Education, Yunnan Normal University, Kunming, China

**Keywords:** tendon, tendinopathy, transcriptomics, pseudobulk, gene-set scoring, reproducibility

## Abstract

We analysed five predefined programmes across three human tendon transcriptomic contexts: acute exercise-associated fibroblast pseudobulks (E-MTAB-15400; 4 control and 4 exercise samples), mechanically stretched donor-derived cells (GSE150482; 3 normal and 3 tendinopathy donors), and paired lesional versus grossly normal-appearing tendon (GSE26051; 23 donor pairs). Inference was conducted at the sample, donor or donor-pair level. Non-stimulated GSE150482 single-cell RNA sequencing comprised only N1 and D1, whereas stretched single-cell RNA sequencing comprised N1–N3 and D1–D3; the conditions also used Drop-seq and 10X Chromium, respectively. A donor-replicated disease-by-stimulation interaction is therefore not estimable. Within-context Benjamini– Hochberg adjustment supported acute mechanical-response, extracellular-matrix (ECM) and protein-folding differences and paired-lesion collagen, integrin and ECM differences, but no stretched donor-background difference. Exact label permutations, a conservative global 15-test adjustment, leave-one-unit analyses, a Fib1–Fib4 pooled sensitivity and a paired exact Wilcoxon sensitivity qualified these findings. Integrin was fully nested in ECM, while collagen overlapped 52/79 genes with ECM. Collagen fibril organization showed the most directionally consistent matrix-associated pattern across the three non-equivalent contrasts, without establishing a disease-specific loading response or causal mechanism. The analysis demonstrates why public tendon datasets must be integrated by estimand and biological inference unit rather than pooled as interchangeable replications.

## 1. Introduction

Tendons adapt to force through coordinated changes in matrix synthesis, organization and cell–matrix signalling, whereas tendinopathy is associated with altered tissue composition and repair biology [1,2]. Determining whether loading responses are preserved, attenuated or reversed in disease requires donor-replicated loading contrasts in both healthy and diseased groups; it cannot be inferred from unrelated cross-sectional contrasts.

Public transcriptomic datasets provide complementary views of human tendon biology, but an acute in-vivo exercise comparison, a comparison between stretched cells from different donor backgrounds and a paired within-donor lesion comparison estimate different quantities. Treating cells as biological replicates or pooling these contrasts can create misleading precision.

Five programmes were defined from MSigDB v2026.1.Hs [3,4]: response to mechanical stimulus, ECM organization, collagen fibril organization, integrin cell-surface interactions and protein folding. They are non-exhaustive candidates and are structurally dependent rather than independent replications.

## 2. Methods

### 2.1 Study governance

The secondary analysis was registered on OSF on 31 August 2026 before formal programme comparisons began on 1 September 2026. The registration is currently embargoed. The registered plan specified the five programmes, biological inference units, primary tests, within-context multiplicity control, exact label-permutation robustness where computationally possible, and unit-deletion analyses. The source-level description of the GSE150482 resting design was corrected during manuscript verification; the registered analysis itself was already restricted to the six stretched donors.

Gene Ontology and Reactome identifiers were interpreted using their primary resource descriptions [5,6]. Computations used R 4.5.2 [7] and Python 3. All 15 primary estimates were recomputed from saved biological-unit score tables and cross-checked against the manuscript tables. Analyses added after registration are labelled as such. OpenAI Codex (OpenAI) was used to assist with drafting and checking code for selected robustness analyses. All AI-assisted code was reviewed by the authors, executed against saved biological-unit data, and its numerical outputs were cross-checked against source tables. AI tools did not determine dataset eligibility, statistical estimands or scientific conclusions.

### 2.2 Programme definitions and scoring

The five memberships were response to mechanical stimulus (GO:0009612; 222 genes), extracellular matrix organization (Reactome R-HSA-1474244; 321 genes), collagen fibril organization (GO:0030199; 79 genes), integrin cell-surface interactions (Reactome R-HSA-216083; 85 genes) and protein folding (GO:0006457; 235 genes). Exact provenance, source version, retrieval date, identifier namespace, membership hashes and scoreable counts are given in Supplementary Table S1. Genes were ranked within each normalized biological-unit profile and scored as one-sided up-sets with singscore [8]. Within each context, the five raw P values formed one family and were adjusted by Benjamini–Hochberg [9].

Overlap was quantified by intersection, union, Jaccard index and asymmetric containment. A programme was strictly nested only if all members were contained in the comparison set. Programme overlap was treated as structural dependence rather than as independent replication.

### 2.3 Acute exercise-associated fibroblast analysis

E-MTAB-15400 profiled the human muscle–tendon unit 4 h after eccentric resistance exercise or control sampling [10]. The registered primary compartment pooled author-annotated Fib1–Fib3 nuclei within each biological sample (17,273 nuclei; 4 control and 4 exercise samples). These populations correspond to tendon/MTJ-associated fibroblasts and were the clusters with exercise-responsive expression in the source report; Fib4 was a small perineural, possibly muscle-resident population [10]. Counts were aggregated by sample, TMM-normalized with edgeR [11], transformed to log counts per million and programme-scored. Two-sided Welch tests reported exercise-minus-control differences, Hedges’ g and 95% CIs.

In the retained n=8 dataset, exercise status was strongly entangled with sex, and sex-specific or condition-by-sex effects could not be estimated reliably. Tendon source was also imbalanced between groups. No overadjusted multivariable model was fitted. The registered primary analysis therefore estimates an exercise-group-associated difference, not an unconfounded causal exercise effect.

A post-registration all-fibroblast sensitivity pooled Fib1–Fib4 using the same raw matrices, author singlet filter, sample-level pseudobulk, gene definitions, normalization, scoring and Welch-test workflow. It was not used to replace the registered Fib1–Fib3 primary analysis.

### 2.4 Stretched donor-background analysis

GSE150482 profiled cultured tendon progenitor cells from normal and tendinopathy donors [12]. Non-stimulated scRNA-seq was available for one normal donor (N1) and one tendinopathy donor (D1), whereas mechanically stimulated scRNA-seq included N1–N3 and D1–D3. The deposited design therefore lacks donor-replicated matched resting transcriptomes for all six donors. Resting and stretched cells were also profiled using Drop-seq and 10X Chromium, respectively. Consequently, a donor-replicated disease-by-stimulation interaction is not estimable.

The estimable contrast used six stretched-arm donor pseudobulks (3 normal-derived and 3 tendinopathy-derived; 10,660 confident singlets at the registered HTO 0.99 threshold). Two-sided Welch tests reported tendinopathy-derived minus normal-derived differences, Hedges’ g and 95% CIs. Leave-one-donor-out analyses were used as a sensitivity check. Cells were not treated as independent disease-group replicates.

### 2.5 Paired lesion analysis

GSE26051 contains 46 Affymetrix U133 Plus 2.0 arrays from 23 donors, each contributing lesional and paired grossly normal-appearing tendon [13]. Arrays underwent joint RMA processing [14,15], probe-to-symbol mapping and median collapse of multiple probes per symbol. Two-sided paired t tests reported lesional-minus-paired-comparator differences, Hedges’ gz and 95% CIs. Nineteen pairs were site-discordant. Leave-one-pair-out and same-site n=4 results were sensitivities; the same-site result is exploratory and descriptive. A post-registration exact two-sided Wilcoxon signed-rank sensitivity was additionally applied to the five paired programme differences, with Benjamini– Hochberg adjustment across those five sensitivity P values.

### 2.6 Robustness and multiplicity sensitivities

For the registered exact-label-permutation analysis, all 70 allocations of 4 of 8 acute samples and all 20 allocations of 3 of 6 stretched donors were enumerated. The two-sided statistic was the absolute Welch t, and the exact P value was the fraction of allocations with an absolute statistic at least as large as the observed statistic. Resolution is necessarily coarse, especially for six donors. The primary parametric tests were not replaced when conclusions differed.

A post-registration conservative multiplicity sensitivity applied Benjamini–Hochberg adjustment once across all 15 primary raw P values. The registered within-context five-test adjustments remained primary. Leave-one-context-out direction summaries were descriptive and did not pool effects.

### 2.7 Interpretive rule

Preserved, attenuated and reversed were reserved for a donor-replicated disease-by-stimulation interaction. A nonsignificant comparison was not interpreted as equivalence. Hedges’ g and paired Hedges’ gz have different variance standards; Figure 5 therefore uses separate panels and x-scales, and no cross-context pooled effect is reported.

## 3. Results

### 3.1 The disease-by-stimulation interaction is not estimable

The corrected deposited design shows resting N1 and D1 only, versus stretched N1–N3 and D1–D3, with workflow also entangled with stimulation (Figure 1; Table 1). The six stretched donor pseudobulks support a disease-background comparison under stretch, but not donor-replicated loading contrasts in both donor groups. None of the programmes can therefore be classified as a preserved, attenuated or reversed loading response.

**Figure 1.**
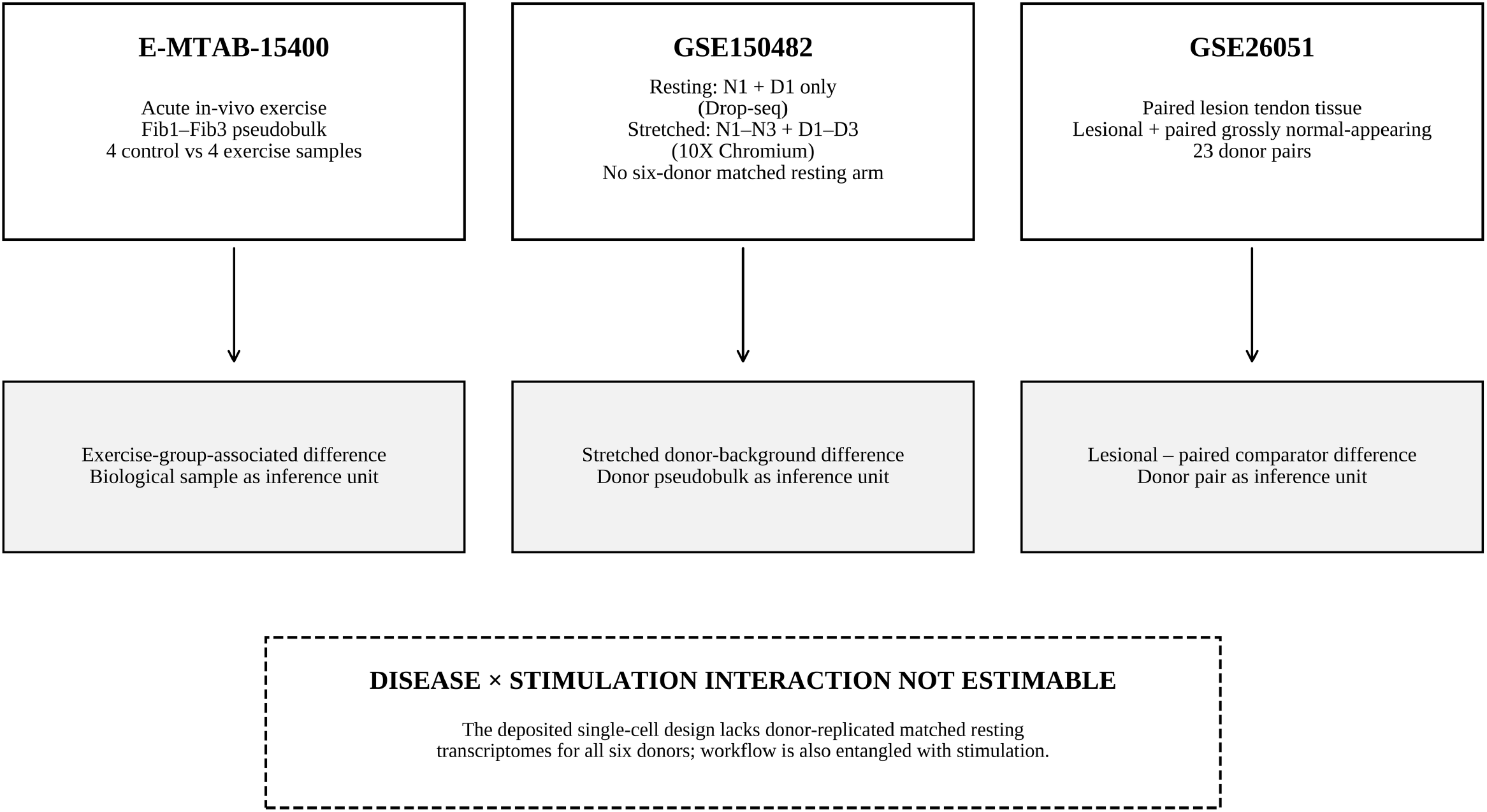
Study design and estimability. Resting GSE150482 scRNA-seq includes N1 and D1 only (Drop-seq); stretched scRNA-seq includes N1–N3 and D1–D3 (10X Chromium). The design lacks a six-donor matched resting arm, so a donor-replicated disease-by-stimulation interaction is not estimable.

**Table 1.** Dataset characteristics, estimands and limitations. Dataset characteristics, estimands and primary limitations.

| Dataset | Material | Contrast | Inference unit | N | Primary limitation |
| --- | --- | --- | --- | --- | --- |
| E-MTAB-15400 | Single nuclei; registered Fib1–Fib3 pseudobulk | Exercise-group-associated vs control | Biological sample | 4 vs 4 | Strong condition–sex imbalance and unequal tendon-source composition. |
| GSE150482 | Cultured tendon progenitor cells | Stretched tendinopathy-derived vs stretched normal-derived | Donor pseudobulk | 3 vs 3 | Resting: N1 and D1 only (Drop-seq); stretched: N1–N3 and D1–D3 (10X); interaction not estimable. |
| GSE26051 | Bulk tendon tissue | Lesional minus paired grossly normal-appearing tendon | Within-donor pair | 23 pairs | 19/23 pairs site-discordant; bulk-composition ambiguity. |

*Non-lesional denotes paired grossly normal-appearing tendon, not tissue from healthy volunteers.*

### 3.2 Primary context-specific estimates

All five acute estimates were positive (Figure 2; Table 2). Mechanical stimulus response was largest (g=5.888, 95% CI 2.420 to 9.328; q=0.000746), followed by protein folding (g=4.988, 1.971 to 7.963; q=0.00760) and ECM organization (g=2.077, 0.373 to 3.699; q=0.0296). Integrin interactions (g=1.634; q=0.0519) and collagen fibril organization (g=0.861; q=0.220) were positive but did not cross the within-context threshold. With four samples per group, standardized effects are unstable [16], and the design precludes exercise-specific attribution.

**Figure 2.**
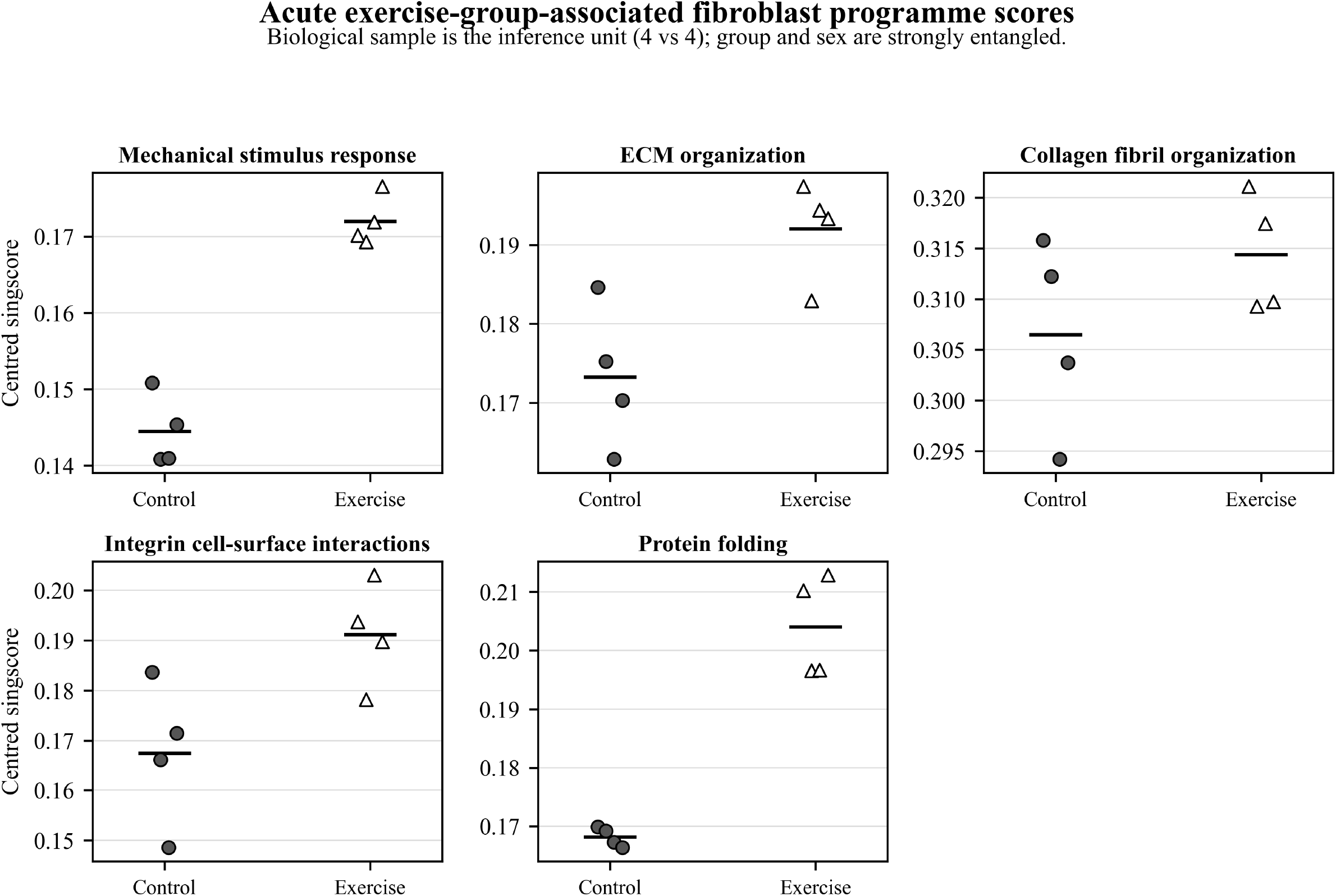
Acute exercise-group-associated programme scores. Each point is one E-MTAB-15400 biological sample. Group and sex are strongly entangled and tendon-source composition is unequal.

**Table 2.** Programme-level estimates. All 15 primary programme estimates recomputed from full-precision biological-unit score tables.

| Context | Programme | N | Effect [95% CI] | P | Within-context q | Direction |
| --- | --- | --- | --- | --- | --- | --- |
| Acute exercise — E-MTAB-15400 | Mechanical response | 4 vs 4 biological samples | 5.888 [2.420, 9.328] | 0.000149 | 0.000746 | positive |
| Stretched donor background — GSE150482 | Mechanical response | 3 vs 3 donors | -0.878 [-2.241, 0.570] | 0.269 | 0.672 | negative |
| Paired lesion — GSE26051 | Mechanical response | 23 donor pairs | 0.300 [-0.107, 0.702] | 0.15 | 0.187 | positive |
| Acute exercise — E-MTAB-15400 | ECM organization | 4 vs 4 biological samples | 2.077 [0.373, 3.699] | 0.0178 | 0.0296 | positive |
| Stretched donor background — GSE150482 | ECM organization | 3 vs 3 donors | -0.044 [-1.319, 1.235] | 0.949 | 0.949 | negative |
| Paired lesion — GSE26051 | ECM organization | 23 donor pairs | 0.475 [0.052, 0.889] | 0.0276 | 0.046 | positive |
| Acute exercise — E-MTAB-15400 | Collagen fibril organization | 4 vs 4 biological samples | 0.861 [-0.463, 2.126] | 0.22 | 0.22 | positive |
| Stretched donor background — GSE150482 | Collagen fibril organization | 3 vs 3 donors | 1.318 [-0.284, 2.818] | 0.126 | 0.632 | positive |
| Paired lesion — GSE26051 | Collagen fibril organization | 23 donor pairs | 0.657 [0.212, 1.090] | 0.00356 | 0.0178 | positive |
| Acute exercise — E-MTAB-15400 | Integrin interactions | 4 vs 4 biological samples | 1.634 [0.088, 3.097] | 0.0415 | 0.0519 | positive |
| Stretched donor background — GSE150482 | Integrin interactions | 3 vs 3 donors | 0.140 [-1.149, 1.412] | 0.846 | 0.949 | positive |
| Paired lesion — GSE26051 | Integrin interactions | 23 donor pairs | 0.541 [0.110, 0.961] | 0.0135 | 0.0338 | positive |
| Acute exercise — E-MTAB-15400 | Protein folding | 4 vs 4 biological samples | 4.988 [1.971, 7.963] | 0.00304 | 0.0076 | positive |
| Stretched donor background — GSE150482 | Protein folding | 3 vs 3 donors | 0.552 [-0.808, 1.850] | 0.446 | 0.743 | positive |
| Paired lesion — GSE26051 | Protein folding | 23 donor pairs | 0.001 [-0.394, 0.395] | 0.996 | 0.996 | positive |

Effects are Hedges' g for unpaired contexts and paired Hedges' gz for GSE26051. Positive directions are exercise higher, tendinopathy-derived higher under stretch, and lesion higher. Direction reports the sign of the point estimate; the leave-one-context descriptive analysis treats |standardized effect|<0.05 as approximately null.

**Table 3.** Estimand-aware cross-context summary. Estimand-aware cross-context summary; no pooled effect is calculated.

| **Programme** | **Acute g [95% CI]** | **Stretched g [95% CI]** | **Paired lesion gz [95% CI]** | **Interpretive boundary** |
| --- | --- | --- | --- | --- |
| Mechanical response | 5.89 [2.42, 9.33] | -0.88 [-2.24, 0.57] | 0.30 [-0.11, 0.70] | Disease × stimulation<br>interaction not estimable |
| ECM organization | 2.08 [0.37, 3.70] | -0.04 [-1.32, 1.24] | 0.48 [0.05, 0.89] | Disease × stimulation<br>interaction not estimable |
| Collagen fibril organization | 0.86 [-0.46, 2.13] | 1.32 [-0.28, 2.82] | 0.66 [0.21, 1.09] | Disease × stimulation<br>interaction not estimable |
| Integrin interactions | 1.63 [0.09, 3.10] | 0.14 [-1.15, 1.41] | 0.54 [0.11, 0.96] | Disease × stimulation<br>interaction not estimable |
| Protein folding | 4.99 [1.97, 7.96] | 0.55 [-0.81, 1.85] | 0.00 [-0.39, 0.40] | Disease × stimulation<br>interaction not estimable |

No stretched donor-background programme reached q<0.05 (Figure 3; Table 2). Estimates were collagen g=1.318 (95% CI −0.284 to 2.818; q=0.632), mechanical response g=-0.878 (−2.241 to 0.570; q=0.672), ECM g=-0.044 (−1.319 to 1.235; q=0.949), integrin g=0.140 (−1.149 to 1.412; q=0.949) and protein folding g=0.552 (−0.808 to 1.850; q=0.743). These describe donor backgrounds after stretch.

**Figure 3.**
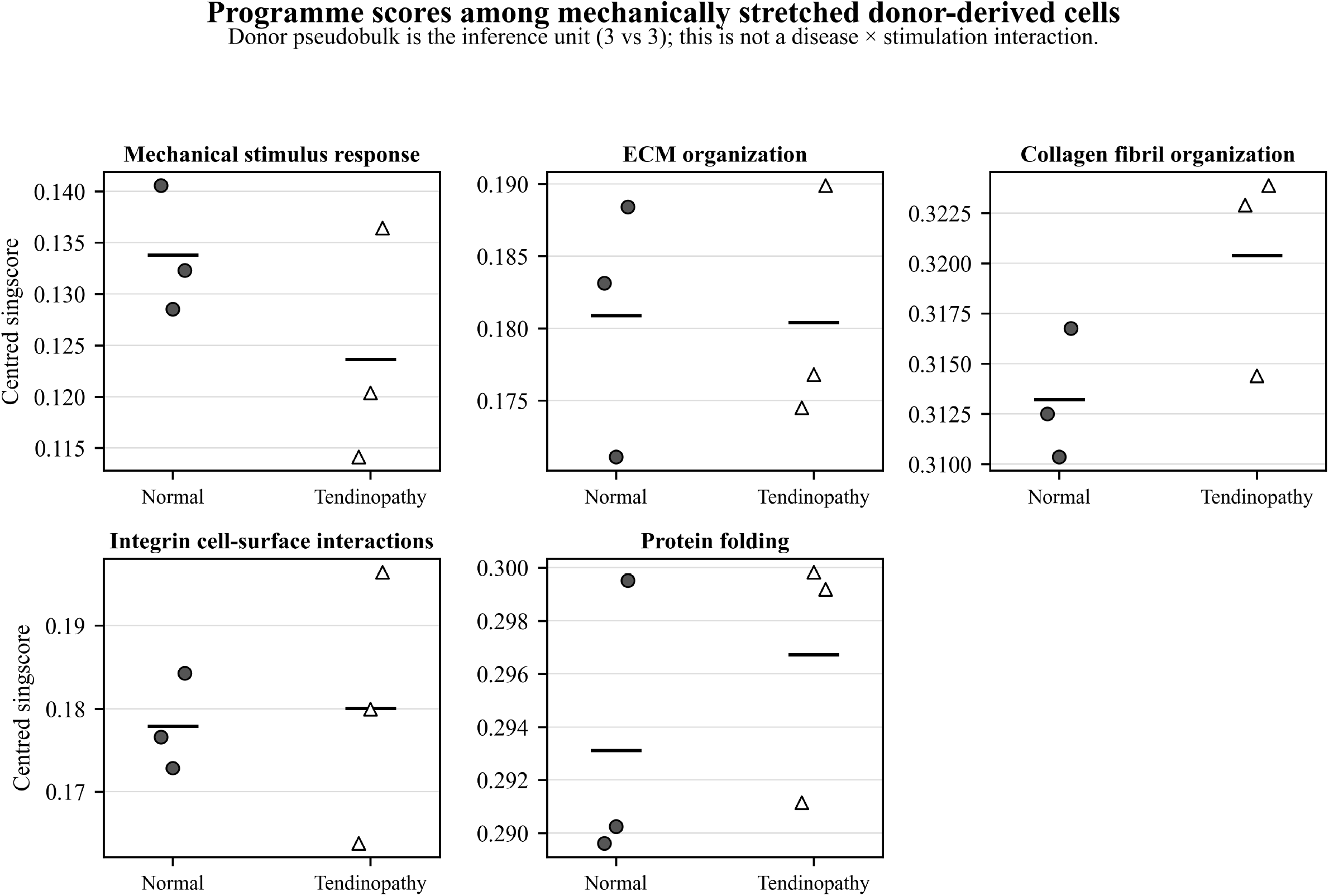
Programme scores among mechanically stretched donor-derived cells. Each point is one GSE150482 donor pseudobulk at HTO threshold 0.99. The contrast is donor background under stretch, not a disease-by-stimulation interaction.

In the paired lesion analysis (Figure 4; Table 2), collagen (gz=0.657, 95% CI 0.212 to 1.090; q=0.0178), integrin (gz=0.541, 0.110 to 0.961; q=0.0338) and ECM (gz=0.475, 0.052 to 0.889; q=0.0460) were higher in lesions. Mechanical response was uncertain (gz=0.300; q=0.187), and protein folding was approximately null (gz=0.001; q=0.996). Bulk-tissue effects combine regulation and composition.

**Figure 4.**
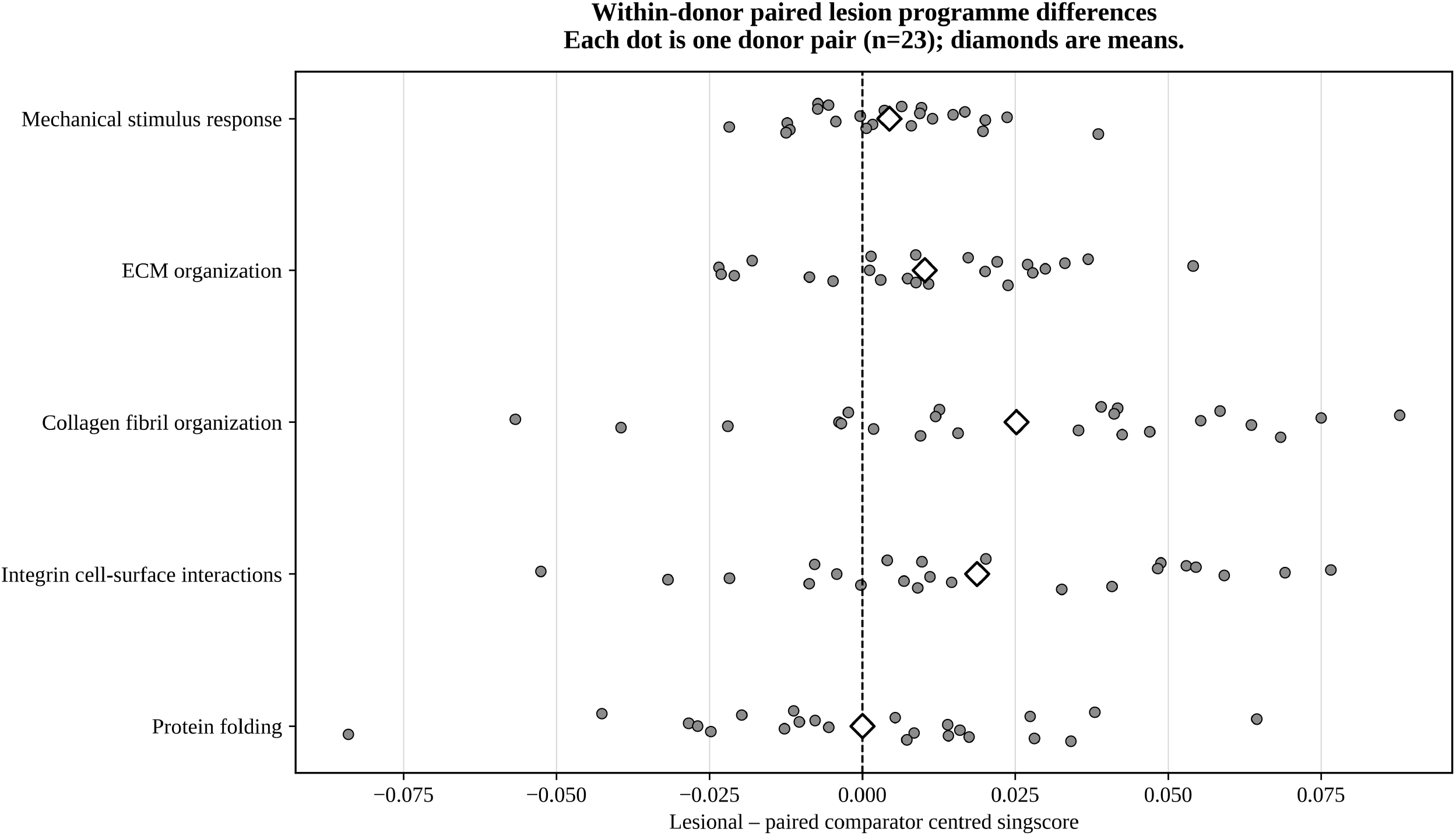
Within-donor paired lesion programme differences. Each point is the lesional-minus-paired-comparator score difference for one of 23 GSE26051 donor pairs; diamonds are means. The paired comparator is grossly normal-appearing tendon from the same donor; 19 pairs are site-discordant.

### 3.3 Programme dependence and cross-context direction

All 85 integrin genes were contained in the 321-gene ECM programme (Jaccard 0.265). ECM contained 52 of 79 collagen genes (65.8%; Jaccard 0.149), and collagen shared 17 genes with integrin (Jaccard 0.116). Integrin and ECM results are therefore not independent. Collagen was positive across all three non-equivalent contrasts and remained positive after any one context was omitted (Figure 5; Supplementary Figures S1 and S2).

**Figure 5.**
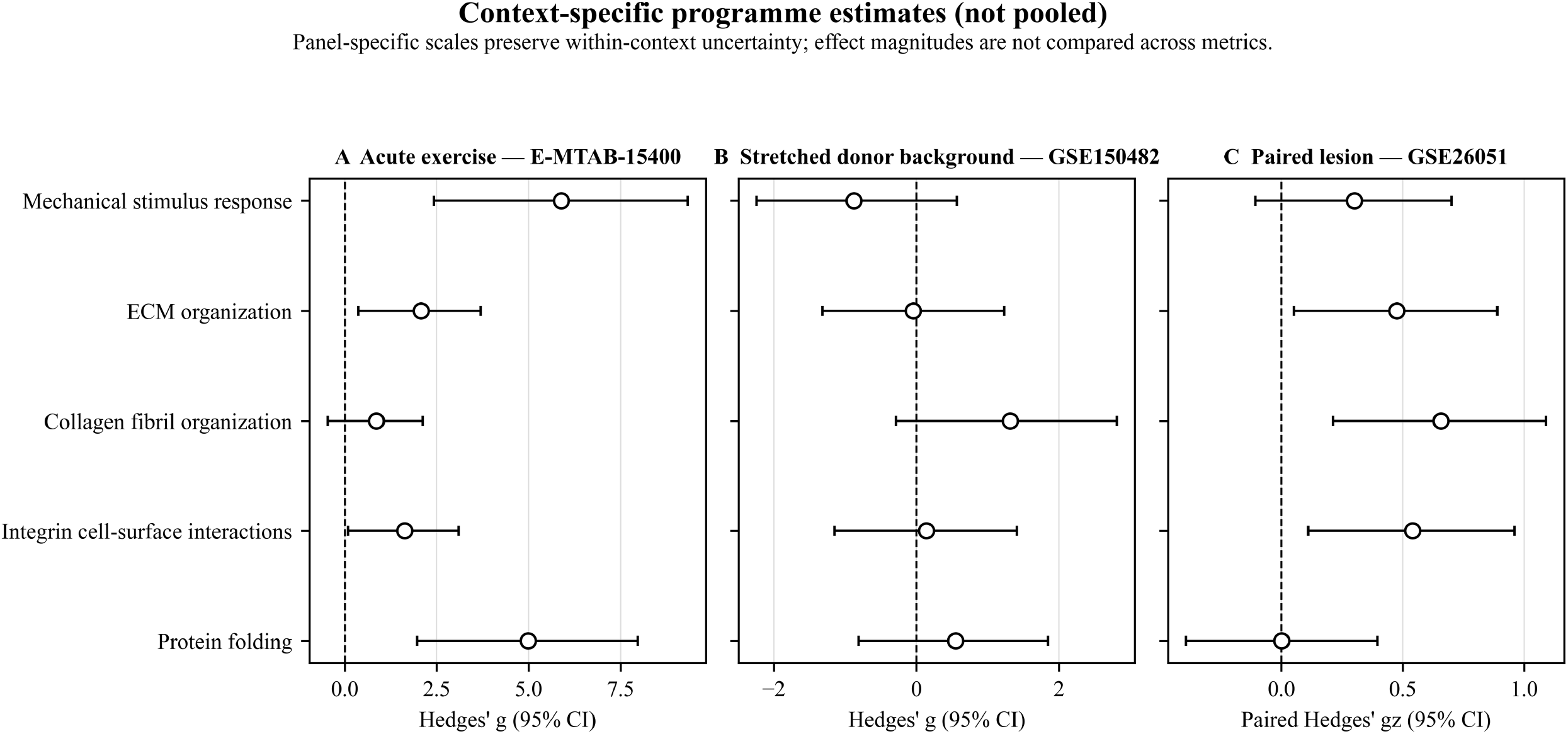
Context-specific estimates without pooling. Panels show Hedges’ g for the two unpaired comparisons and paired Hedges’ gz for the paired-lesion contrast. Each panel has its own scale; magnitudes are not compared across metrics.

### 3.4 Robustness results

The Fib1–Fib4 pooled sensitivity retained positive directions for all five programmes. Its Hedges’ g estimates were 7.754 (mechanical response), 1.950 (ECM), 0.754 (collagen), 1.309 (integrin) and 4.951 (protein folding); within-sensitivity q values were 0.000129, 0.0411, 0.270, 0.0985 and 0.00614, respectively. This post-registration analysis supports directional robustness to including the small Fib4 population but does not resolve group confounding.

Exact label-permutation P values in the acute comparison were 0.0286 for mechanical response and protein folding, 0.0571 for ECM and integrin, and 0.257 for collagen. For the six stretched donors, values were 0.300, 1.000, 0.200, 0.900 and 0.300, respectively; the 20-allocation resolution limits calibration. Under global 15-test BH, q values below 0.05 remained for acute mechanical response (0.00224), acute protein folding (0.0178) and paired-lesion collagen (0.0178). In the paired-lesion exact Wilcoxon sensitivity, raw P values (BH q) were 0.200 (0.250) for mechanical response, 0.0254 (0.0424) for ECM, 0.00485 (0.0243) for collagen, 0.0135 (0.0338) for integrin and 0.846 (0.846) for protein folding (Supplementary Table S5).

All 15 estimates recomputed from full-precision biological-unit tables matched the saved primary results within 1 × 10^-10. Acute directions persisted through all eight leave-one-sample analyses. In stretched cells, mechanical response, collagen and protein folding retained direction in all six donor deletions, whereas ECM did so in two and integrin in four. In the paired lesion context, the first four programmes retained direction through all 23 pair deletions; protein folding changed sign in 12. The same-site n=4 subset was positive for the four matrix/mechanical programmes and negative for protein folding, but is exploratory.

## 4. Discussion

The central result is an estimability constraint, not a negative interaction. Because resting scRNA-seq includes only N1 and D1 and uses a different workflow from stretched samples, the deposited design cannot estimate donor-replicated disease-specific loading responses. This distinction prevents a six-donor stretched comparison from being mislabelled as an interaction.

Collagen fibril organization showed the most consistent direction among the matrix-associated programmes and, unlike integrin, was not fully nested within the broader ECM set. Its overlap with ECM remains substantial (52/79 genes), so its recurrence is not independent of the broader matrix domain. The paired-lesion result and leave-one-context-out direction nominate collagen organization for prospective testing, but do not prove preserved loading responsiveness or causality.

The acute separation for mechanical response, ECM and protein folding must be read against strong condition–sex imbalance, unequal tendon-source composition and n=8. The all-fibroblast sensitivity indicates that excluding Fib4 does not determine direction, but adding Fib4 cannot remove design confounding. Similarly, the stretched comparison has only three donors per group and coarse permutation resolution. The paired-lesion analysis preserves donor pairing but most pairs differ by anatomical site and bulk expression cannot distinguish transcriptional regulation from cell composition.

The conservative global BH sensitivity changes the multiplicity question and therefore supplements rather than replaces the registered within-context families. Exact permutations, paired nonparametric sensitivity and unit-deletion analyses calibrate confidence without pooling non-equivalent effects.

These programme choices are biologically plausible because tendon loading engages mechanotransduction, matrix remodelling, collagen synthesis, integrin–ILK–AKT–mTOR signalling and proteostasis [17–27]. The published β1-integrin/ILK/AKT/mTOR/collagen evidence specifically supports pathway rationale rather than a causal interpretation of these observational reanalyses [21].

A decisive study would sample the same healthy and tendinopathy donors before and after standardized loading, balance sex, record tendon source and use a donor-level disease-by-time interaction with a prespecified equivalence margin. Cell-resolved profiling could then localize, rather than replace, the donor-level interaction.

## 5. Conclusions

The recoverable data do not identify whether tendinopathy preserves, attenuates or reverses an acute loading response. Collagen fibril organization showed the most consistent recurrent matrix-associated direction, but its substantial overlap with the broader ECM programme cautions against interpreting it as an independent mechanism. The principal contribution is an estimand-aware, donor-aware and programme-dependency-aware framework that preserves biological inference units and avoids pooling non-equivalent public datasets.

## Data availability

The source datasets are public: E-MTAB-15400 (https://www.ebi.ac.uk/biostudies/studies/E-MTAB-15400), GSE150482 (https://www.ncbi.nlm.nih.gov/geo/query/acc.cgi?acc=GSE150482) and GSE26051 (https://www.ncbi.nlm.nih.gov/geo/query/acc.cgi?acc=GSE26051). No new participant-level data were generated. The OSF registration remains embargoed. Derived biological-unit score tables and analysis code are supplied in the accompanying reproducibility archive.

## Code availability

The accompanying reproducibility archive contains statistical reproduction scripts, biological-unit source tables, programme definitions, figure source data, software information, execution order and SHA-256 manifests. It does not redistribute restricted or identifying data. No external repository or DOI was created during this work.

## Author contributions

Yushuo Liu: Conceptualization, Data curation, Formal analysis, Methodology, Software, Validation, Visualization, Writing – original draft, Writing – review & editing. Zitao Wang: Validation, Writing – review & editing. Qiuyuan Peng: Validation, Writing – review & editing. Yang Li: Validation, Writing – review & editing. Bingao Chen: Supervision, Project administration, Methodology, Writing – review & editing.

## Ethics

This study is a secondary analysis of publicly available, de-identified transcriptomic datasets. No new participants or specimens were recruited and no identifiable participant information was accessed. Ethics approval and consent for the original data collections are described in the source studies [10,12,13].

## Declaration of generative AI and AI-assisted technologies in the manuscript preparation process

During the preparation of this work, the authors used OpenAI ChatGPT and Codex to improve language and readability and to assist with manuscript organization, pre-submission consistency checking, and document and figure/table layout quality control. OpenAI Codex was also used as described in Methods to assist with drafting and checking code for selected robustness analyses. After using these tools, the authors reviewed and edited the content and code as needed and take full responsibility for the content of the article. No generative AI was used to fabricate, replace or alter the underlying research data.

