## Supplementary Materials for "Estimand-aware and donor-aware triangulation of predefined gene-set signals across human tendon transcriptomic datasets"

**Supplementary Methods**

The supplementary analyses used the same biological inference units, fixed programme memberships and rank-based scoring as the main analysis. Exact label permutations were specified in the registered plan. The Fib1–Fib4 pooled analysis, paired exact Wilcoxon signed-rank analysis and global 15-test Benjamini–Hochberg adjustment were post-registration sensitivities.

**Supplementary Figure Legends**

Supplementary Figure S1. Pairwise programme overlap. Grayscale cells report Jaccard indices; complete asymmetric containment is in Table S2.

Supplementary Figure S2. Leave-one-context-out directional robustness. Each cell shows directions in the two retained non-equivalent contrasts; 0 denotes an absolute standardized effect below 0.05. No pooling.

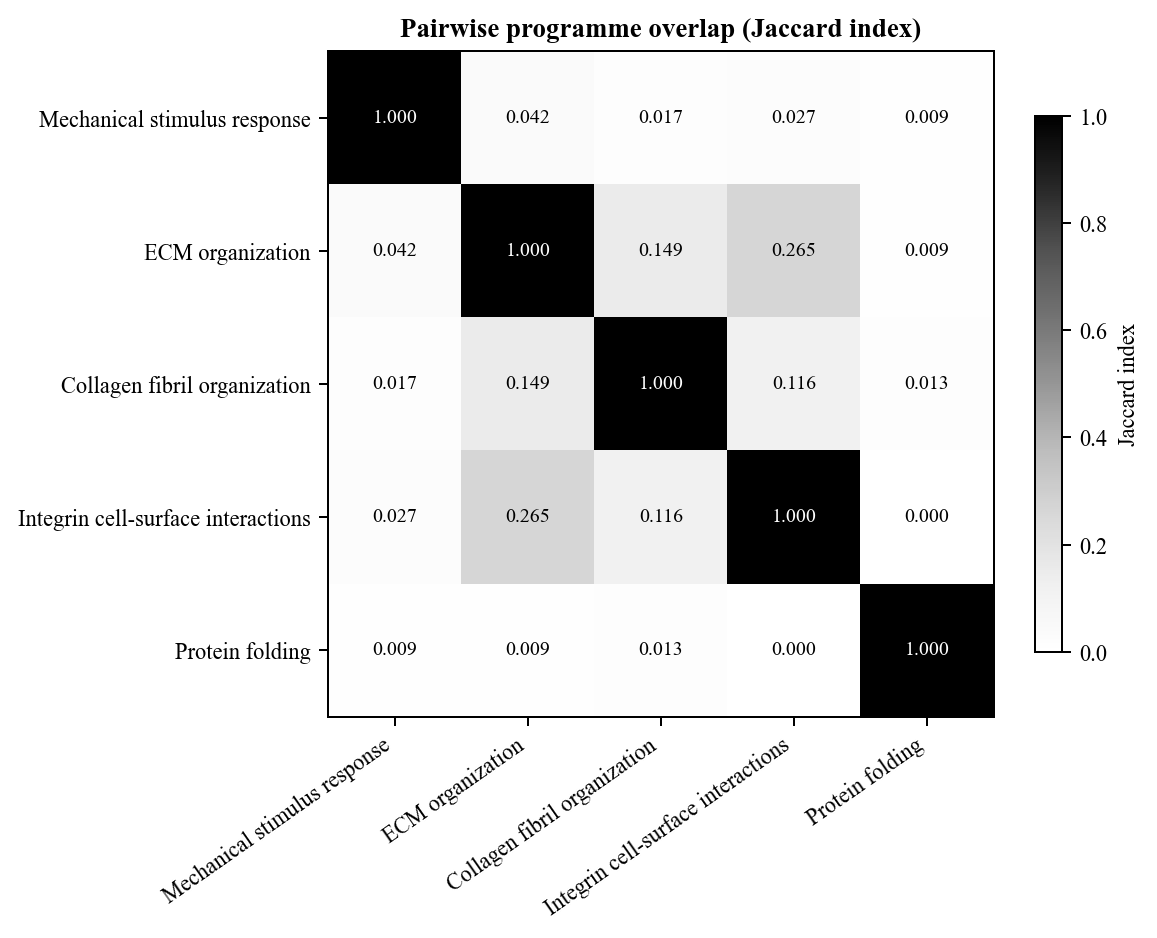

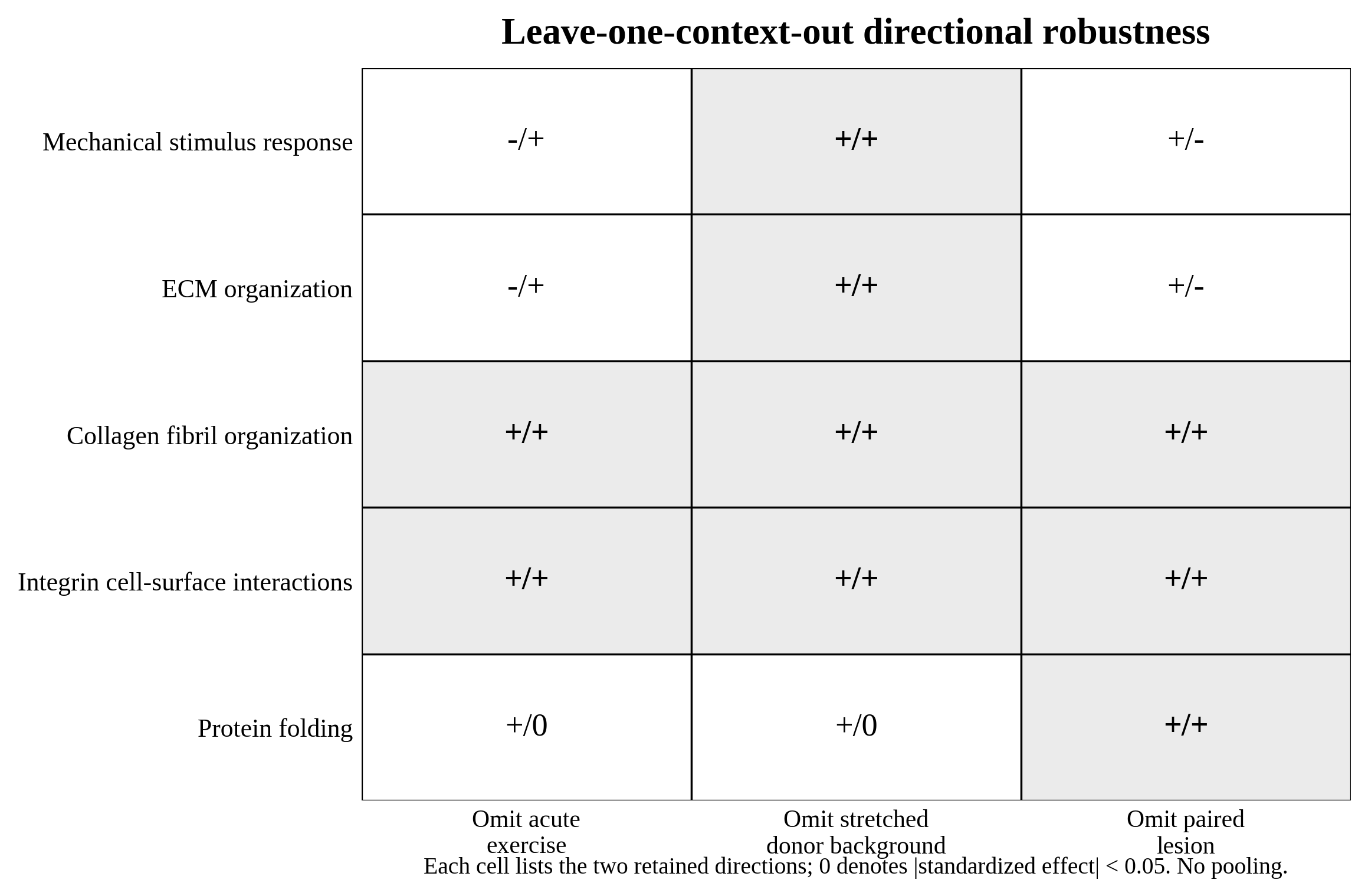

**Table S1. Programme definition provenance and scoreable counts.**

| **Exact name** | **Source** | **ID** | **Version** | **MSigDB ID** | **Retrieved** | **Original n** | **Scoreable n: E-MTAB/GSE150482/GSE26051** | **Namespace** | **Definition file** | **SHA-256** |
| --- | --- | --- | --- | --- | --- | --- | --- | --- | --- | --- |
| GOBP_RESPONSE_TO_MECHANICAL_STIMULUS | GO | GO:0009612 | v2026.1.Hs | M16378 | 2026-08-31 | 222 | 221/209/212 | HGNC symbols | No aggregate GMT retained; exact per-set TSV in archive | aa768eabb0ca1049a15d6e3e28645cc8ea5d294afcddd3e6b383fbbc9745e2fd |
| REACTOME_EXTRACELLULAR_MATRIX_ORGANIZATION | REACTOME | R-HSA-1474244 | v2026.1.Hs | M610 | 2026-08-31 | 321 | 321/314/318 | HGNC symbols | No aggregate GMT retained; exact per-set TSV in archive | 0bec96565c2a1b5e40c07d3d2689859a43daab731b11b01ce4e42b846dabf92e |
| GOBP_COLLAGEN_FIBRIL_ORGANIZATION | GO | GO:0030199 | v2026.1.Hs | M10505 | 2026-08-31 | 79 | 78/73/74 | HGNC symbols | No aggregate GMT retained; exact per-set TSV in archive | b7c00a97baa8a124e9560462a8a0a242b9f091c34aa22ad79a897c6ffa2306e4 |
| REACTOME_INTEGRIN_CELL_SURFACE_INTERACTIONS | REACTOME | R-HSA-216083 | v2026.1.Hs | M16441 | 2026-08-31 | 85 | 85/84/85 | HGNC symbols | No aggregate GMT retained; exact per-set TSV in archive | 9437a834febaadb429f8753bb8dbe57c8970c3c94af5128732c07f02bdf205f5 |
| GOBP_PROTEIN_FOLDING | GO | GO:0006457 | v2026.1.Hs | M13856 | 2026-08-31 | 235 | 224/215/213 | HGNC symbols | No aggregate GMT retained; exact per-set TSV in archive | e81134dbd3d2ce09e6458d6da3ccbc97379424912c3b931c507dd178da0e318b |

**Table S2. Ordered-pair programme overlap and containment.**

| **Set A** | **Set B** | **nA** | **nB** | **Shared** | **Jaccard** | **A in B** | **B in A** | **Nesting** |
| --- | --- | --- | --- | --- | --- | --- | --- | --- |
| Mechanical response | Mechanical response | 222 | 222 | 222 | 1.0000 | 1.0000 | 1.0000 | Identical |
| Mechanical response | ECM organization | 222 | 321 | 22 | 0.0422 | 0.0991 | 0.0685 | Not nested |
| Mechanical response | Collagen fibril organization | 222 | 79 | 5 | 0.0169 | 0.0225 | 0.0633 | Not nested |
| Mechanical response | Integrin interactions | 222 | 85 | 8 | 0.0268 | 0.0360 | 0.0941 | Not nested |
| Mechanical response | Protein folding | 222 | 235 | 4 | 0.0088 | 0.0180 | 0.0170 | Not nested |
| ECM organization | Mechanical response | 321 | 222 | 22 | 0.0422 | 0.0685 | 0.0991 | Not nested |
| ECM organization | ECM organization | 321 | 321 | 321 | 1.0000 | 1.0000 | 1.0000 | Identical |
| ECM organization | Collagen fibril organization | 321 | 79 | 52 | 0.1494 | 0.1620 | 0.6582 | Not nested |
| ECM organization | Integrin interactions | 321 | 85 | 85 | 0.2648 | 0.2648 | 1.0000 | Set B is a strict subset of Set A |
| ECM organization | Protein folding | 321 | 235 | 5 | 0.0091 | 0.0156 | 0.0213 | Not nested |
| Collagen fibril organization | Mechanical response | 79 | 222 | 5 | 0.0169 | 0.0633 | 0.0225 | Not nested |
| Collagen fibril organization | ECM organization | 79 | 321 | 52 | 0.1494 | 0.6582 | 0.1620 | Not nested |
| Collagen fibril organization | Collagen fibril organization | 79 | 79 | 79 | 1.0000 | 1.0000 | 1.0000 | Identical |
| Collagen fibril organization | Integrin interactions | 79 | 85 | 17 | 0.1156 | 0.2152 | 0.2000 | Not nested |
| Collagen fibril organization | Protein folding | 79 | 235 | 4 | 0.0129 | 0.0506 | 0.0170 | Not nested |
| Integrin interactions | Mechanical response | 85 | 222 | 8 | 0.0268 | 0.0941 | 0.0360 | Not nested |
| Integrin interactions | ECM organization | 85 | 321 | 85 | 0.2648 | 1.0000 | 0.2648 | Set A is a strict subset of Set B |
| Integrin interactions | Collagen fibril organization | 85 | 79 | 17 | 0.1156 | 0.2000 | 0.2152 | Not nested |
| Integrin interactions | Integrin interactions | 85 | 85 | 85 | 1.0000 | 1.0000 | 1.0000 | Identical |
| Integrin interactions | Protein folding | 85 | 235 | 0 | 0.0000 | 0.0000 | 0.0000 | Not nested |
| Protein folding | Mechanical response | 235 | 222 | 4 | 0.0088 | 0.0170 | 0.0180 | Not nested |
| Protein folding | ECM organization | 235 | 321 | 5 | 0.0091 | 0.0213 | 0.0156 | Not nested |
| Protein folding | Collagen fibril organization | 235 | 79 | 4 | 0.0129 | 0.0170 | 0.0506 | Not nested |
| Protein folding | Integrin interactions | 235 | 85 | 0 | 0.0000 | 0.0000 | 0.0000 | Not nested |
| Protein folding | Protein folding | 235 | 235 | 235 | 1.0000 | 1.0000 | 1.0000 | Identical |

**Table S3. Leave-one-context-out descriptive robustness.**

| **Programme** | **Omitted** | **Retained 1** | **Retained 2** | **Direction 1** | **Direction 2** | **Summary** |
| --- | --- | --- | --- | --- | --- | --- |
| Mechanical response | Acute exercise | Stretched donor background | Paired lesion | - | + | mixed directions |
| Mechanical response | Stretched donor background | Acute exercise | Paired lesion | + | + | same positive direction |
| Mechanical response | Paired lesion | Acute exercise | Stretched donor background | + | - | mixed directions |
| ECM organization | Acute exercise | Stretched donor background | Paired lesion | - | + | mixed directions |
| ECM organization | Stretched donor background | Acute exercise | Paired lesion | + | + | same positive direction |
| ECM organization | Paired lesion | Acute exercise | Stretched donor background | + | - | mixed directions |
| Collagen fibril organization | Acute exercise | Stretched donor background | Paired lesion | + | + | same positive direction |
| Collagen fibril organization | Stretched donor background | Acute exercise | Paired lesion | + | + | same positive direction |
| Collagen fibril organization | Paired lesion | Acute exercise | Stretched donor background | + | + | same positive direction |
| Integrin interactions | Acute exercise | Stretched donor background | Paired lesion | + | + | same positive direction |
| Integrin interactions | Stretched donor background | Acute exercise | Paired lesion | + | + | same positive direction |
| Integrin interactions | Paired lesion | Acute exercise | Stretched donor background | + | + | same positive direction |
| Protein folding | Acute exercise | Stretched donor background | Paired lesion | + | 0 | positive / approximately null |
| Protein folding | Stretched donor background | Acute exercise | Paired lesion | + | 0 | positive / approximately null |
| Protein folding | Paired lesion | Acute exercise | Stretched donor background | + | + | same positive direction |

**Table S4. Biological-unit and all-fibroblast sensitivities.**

| **Context** | **Sensitivity** | **Programme** | **Result** | **Interpretation** |
| --- | --- | --- | --- | --- |
| Acute exercise — E-MTAB-15400 | Fib1–Fib4 pooled all-fibroblast | Mechanical response | g=7.754 [3.325, 12.179]; P=2.57e-05; q=0.000129 | Post-registration; all directions exercise higher |
| Acute exercise — E-MTAB-15400 | Fib1–Fib4 pooled all-fibroblast | ECM organization | g=1.950 [0.293, 3.524]; P=0.0246; q=0.0411 | Post-registration; all directions exercise higher |
| Acute exercise — E-MTAB-15400 | Fib1–Fib4 pooled all-fibroblast | Collagen fibril organization | g=0.754 [-0.547, 2.001]; P=0.27; q=0.27 | Post-registration; all directions exercise higher |
| Acute exercise — E-MTAB-15400 | Fib1–Fib4 pooled all-fibroblast | Integrin interactions | g=1.309 [-0.134, 2.675]; P=0.0788; q=0.0985 | Post-registration; all directions exercise higher |
| Acute exercise — E-MTAB-15400 | Fib1–Fib4 pooled all-fibroblast | Protein folding | g=4.951 [1.953, 7.909]; P=0.00246; q=0.00614 | Post-registration; all directions exercise higher |
| Acute exercise — E-MTAB-15400 | Leave one biological sample out | All programmes | Directions retained in all 8 deletions | Does not resolve confounding |
| Stretched donor background — GSE150482 | Leave one donor out | All programmes | Direction retained: mechanical 6/6; ECM 2/6; collagen 6/6; integrin 4/6; protein folding 6/6 | ECM and integrin donor-sensitive |
| Paired lesion — GSE26051 | Same-site subset | All programmes | n=4; first four positive, protein folding negative | Exploratory/descriptive only |

**Table S5. Paired-lesion exact Wilcoxon signed-rank sensitivity.**

| **Programme** | **Pairs** | **Wilcoxon W** | **Exact two-sided P** | **BH q** | **Interpretation** |
| --- | --- | --- | --- | --- | --- |
| Mechanical stimulus response | 23 | 95 | 0.2002 | 0.2502 | Post-registration nonparametric sensitivity |
| Extracellular matrix organization | 23 | 65 | 0.02542 | 0.04237 | Post-registration nonparametric sensitivity |
| Collagen fibril organization | 23 | 48 | 0.004851 | 0.02426 | Post-registration nonparametric sensitivity |
| Integrin cell-surface interactions | 23 | 58 | 0.01353 | 0.03382 | Post-registration nonparametric sensitivity |
| Protein folding | 23 | 131 | 0.8462 | 0.8462 | Post-registration nonparametric sensitivity |

**Table S6. Registered exact label-permutation robustness.**

| **Context** | **Programme** | **Units** | **Allocations** | **\|Tobs\|** | **Extreme** | **Exact two-sided P** |
| --- | --- | --- | --- | --- | --- | --- |
| Acute exercise — E-MTAB-15400 | Mechanical stimulus response | 8 | 70 | 9.5860 | 2 | 0.0286 |
| Acute exercise — E-MTAB-15400 | Extracellular matrix organization | 8 | 70 | 3.3822 | 4 | 0.0571 |
| Acute exercise — E-MTAB-15400 | Collagen fibril organization | 8 | 70 | 1.4021 | 18 | 0.2571 |
| Acute exercise — E-MTAB-15400 | Integrin cell-surface interactions | 8 | 70 | 2.6595 | 4 | 0.0571 |
| Acute exercise — E-MTAB-15400 | Protein folding | 8 | 70 | 8.1202 | 2 | 0.0286 |
| Stretched donor background — GSE150482 | Mechanical stimulus response | 6 | 20 | 1.3482 | 6 | 0.3000 |
| Stretched donor background — GSE150482 | Extracellular matrix organization | 6 | 20 | 0.0682 | 20 | 1.0000 |
| Stretched donor background — GSE150482 | Collagen fibril organization | 6 | 20 | 2.0229 | 4 | 0.2000 |
| Stretched donor background — GSE150482 | Integrin cell-surface interactions | 6 | 20 | 0.2149 | 18 | 0.9000 |
| Stretched donor background — GSE150482 | Protein folding | 6 | 20 | 0.8466 | 6 | 0.3000 |

**Table S7. Conservative global 15-test multiplicity sensitivity.**

| **Context** | **Programme** | **Raw P** | **Primary within-context q** | **Global 15-test q** |
| --- | --- | --- | --- | --- |
| Acute exercise — E-MTAB-15400 | Mechanical stimulus response | 0.00014915 | 0.000745751 | 0.00223725 |
| Stretched donor background — GSE150482 | Mechanical stimulus response | 0.268715 | 0.671788 | 0.36643 |
| Paired lesion — GSE26051 | Mechanical stimulus response | 0.149737 | 0.187172 | 0.249562 |
| Acute exercise — E-MTAB-15400 | Extracellular matrix organization | 0.0177769 | 0.0296281 | 0.0533307 |
| Stretched donor background — GSE150482 | Extracellular matrix organization | 0.948923 | 0.948923 | 0.996436 |
| Paired lesion — GSE26051 | Extracellular matrix organization | 0.0275795 | 0.0459658 | 0.0689487 |
| Acute exercise — E-MTAB-15400 | Collagen fibril organization | 0.220499 | 0.220499 | 0.330748 |
| Stretched donor background — GSE150482 | Collagen fibril organization | 0.126447 | 0.632236 | 0.237089 |
| Paired lesion — GSE26051 | Collagen fibril organization | 0.00356119 | 0.017806 | 0.017806 |
| Acute exercise — E-MTAB-15400 | Integrin cell-surface interactions | 0.0415222 | 0.0519028 | 0.0889762 |
| Stretched donor background — GSE150482 | Integrin cell-surface interactions | 0.846109 | 0.948923 | 0.976279 |
| Paired lesion — GSE26051 | Integrin cell-surface interactions | 0.0135031 | 0.0337577 | 0.0506365 |
| Acute exercise — E-MTAB-15400 | Protein folding | 0.00303882 | 0.00759704 | 0.017806 |
| Stretched donor background — GSE150482 | Protein folding | 0.445734 | 0.74289 | 0.557168 |
| Paired lesion — GSE26051 | Protein folding | 0.996436 | 0.996436 | 0.996436 |

**Table S8. Dataset eligibility and estimability decisions.**

| **Accession** | **Decision** | **Potential role** | **Inference unit** | **Reason/limitation** |
| --- | --- | --- | --- | --- |
| E-MTAB-15400 | Included | Acute exercise-associated fibroblast pseudobulk | 4 vs 4 biological samples | Strong condition–sex imbalance; unequal tendon-source composition |
| GSE150482 | Partially included | Disease background among stretched donor pseudobulks | 3 vs 3 donors | Resting N1 and D1 only; different workflows; interaction not estimable |
| GSE26051 | Included | Lesional minus paired grossly normal-appearing tendon | 23 donor pairs | 19/23 site-discordant; bulk composition |
| E-MTAB-12530 | Not analysed for localization | Candidate spatial localization resource | Not taken forward | Available object lacked direct machine-readable labels required by the registered gate |
| GSE293788 | Not added | Potential disease-background dataset | Unpaired healthy vs tendinopathy | Not equivalent to paired lesion estimand; no valid pooling |
